# ANDROMEDA by Prosilico and log D outperform human hepatocytes for the prediction of intrinsic hepatic metabolic clearance of carboxylic acids

**DOI:** 10.1101/2023.12.04.569912

**Authors:** Urban Fagerholm

## Abstract

**Introduction:** Extrahepatic metabolism/conjugation, deconjugation of their metabolites, and low and varying unbound fraction in plasma (f_u_), is characteristic for carboxylic drugs. Thus, it is comparably difficult to estimate their *in vivo* intrinsic hepatic metabolic clearance (CL_int_) and hepatic CL (CL_H_) and to predict their *in vivo* CL_int_, CL_H_ and CL. One objective was to investigate the laboratory variability of f_u_ and CL_int_ for carboxylic acids. Another objective was to compare human hepatocytes, measured log D and the software ANDROMEDA with regards to prediction of human *in vivo* CL_int_ of carboxylic acids.

**Materials and Methods:** Measured unbound hepatocyte CL_int_, non-renal CL (surrogate for CL_H_), non-renal CL_int_ (surrogate for hepatic metabolic CL_int_), log D and f_u_ data were taken from studies in the literature. ANDROMEDA (by Prosilico; version 1.0) prediction software was used for *in silico* predictions of CL_int_ for carboxylic acids not used in the training set of its CL_int_-model.

**Results and Discussion:** Mean and maximum differences between highest and lowest reported *in vivo* CL_int_ predicted from hepatocyte CL_int_ were 210- and 1,476-fold (n=8), respectively. Corresponding estimates for *in vitro* f_u_ were 19- and 50-fold, respectively. The data set with the apparently highest number of carboxylic acids contains 39 carboxylic acids with *in vitro* CL_int_ and log D (both measured at the same laboratory), *in vivo* CL_int_ and *in vitro* f_u_. 18 carboxylic acids were excluded as their *in vitro* CL_int_ was below the limit of quantification. The correlation coefficient (R^2^) for log hepatocyte predicted *in vivo* CL_int_ *vs* log *in vivo* CL_int_ was 0.34. The corresponding R^2^ for log D *vs* log *in vivo* CL_int_ was 0.40 (0.47 for 64 carboxylic acids). The Q^2^ (forward-looking R^2^) for *in silico* (ANDROMEDA) predicted and measured log *in vivo* CL_int_ for 12 carboxylic acids was 0.86. The corresponding R^2^ for hepatocytes and log D were 0.67 and 0.66, respectively. ANDROMEDA produced a lower maximum prediction error compared to hepatocytes and also predicted the *in vivo* CL_int_ for all carboxylic acids out of reach for the hepatocyte assay.

**Conclusion:** Very large interlaboratory variability was demonstrated for plasma protein binding and hepatocyte assays. Log D, and especially ANDROMEDA, outperformed the hepatocyte assay for the prediction of CL_int_ of carboxylic acids *in vivo* in man.

## Introduction

Extrahepatic metabolism/conjugation is characteristic for carboxylic acid drugs. In addition to conjugation in the liver, the kidneys and intestines are important organs for conjugation of carboxylic drugs (Soars et al. 2002). This makes predictions of their *in vivo* intrinsic hepatic metabolic clearance (CL_int_), hepatic CL (CL_H_) and CL more difficult. Drug-conjugates are often excreted in bile and might undergo deconjugation (back to drug) and reabsorption in the intestines (such as for the carboxylic acid telmisartan), which adds complexity. Many carboxylic acids have low and varying unbound fraction in plasma (f_u_), which further increases the complexity and uncertainty.

One objective of the study was to investigate the laboratory variability of f_u_ and CL_int_ for carboxylic acids. Another objective was to compare human hepatocytes, measured log D and the software ANDROMEDA by Prosilico with regards to prediction of *in vivo* CL_int_ of carboxylic acids.

## Materials & Methods

The literature was searched for studies with large(st) amount of data for measured unbound hepatocyte CL_int_, non-renal CL (surrogate for CL_H_), non-renal CL_int_ (surrogate for hepatic metabolic CL_int_), log D and f_u_. Useful data were found in the following references: Bowman and Benet 2019 (predicted *in vivo* CL_int_ with human hepatocytes; data from several studies), Riley et al. 2005 (predicted *in vivo* CL_int_ with human hepatocytes), Keefer et al. 2023 (predicted *in vivo* CL_int_ with human hepatocytes in their laboratory, measured log D in their laboratory, and *in vivo* CL_int_), Fagerholm et al. 2021 (f_u_-data from many sources) and Sohlenius-Sternbeck et al. 2010 (predicted *in vivo* CL_int_ with human hepatocytes in their laboratory).

ANDROMEDA by Prosilico (version 1.0) prediction software was used for *in silico* predictions of CL_int_ for carboxylic acids not used in the training set of the CL_int_-model.

## Results & Discussion

### Laboratory variability

Eight carboxylic acids with measured f_u_ and predicted *in vivo* CL_int_ data from various sources were found - diclofenac, fenoprofen, gemfibrozil, ibuprofen, indomethacin, ketoprofen, montelukast and naproxen.

Mean and maximum differences between highest and lowest reported *in vivo* CL_int_ for these predicted from hepatocyte CL_int_ were 210- and 1,476-fold, respectively. (Figures 1 and 2). Corresponding estimates for *in vitro* f_u_ were 19- and 50-fold, respectively (Figure 2). Particularly high CL_int_ • f_u_-differences (between highest and lowest) were found for diclofenac (628-fold), fenoprofen (225-fold), gemfibrozil (1,676-fold), ibuprofen (134-fold), montelukast (16,404-fold) and naproxen (138-fold). Thus, quite large interlaboratory variability and low reproducibility was found for these selected carboxylic acids. This is believed to have a significant impact on human pharmacokinetic, exposure and dose predictions. The findings show general limitations with human hepatocytes and plasma protein measurements when predicting the pharmacokinetics of carboxylic acids.

**Figure 1.**
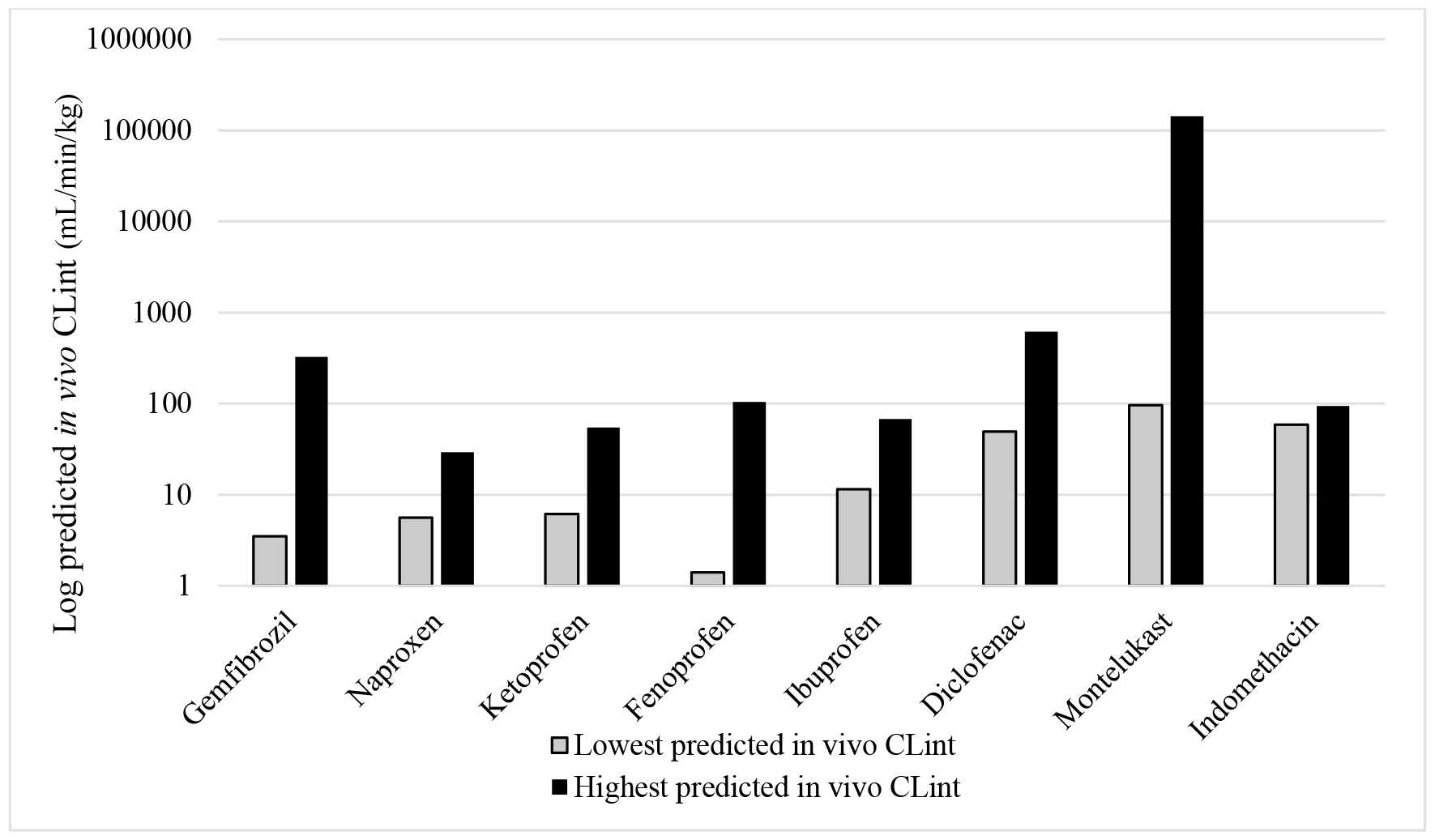
Lowest and highest reported predicted *in vivo* log CL_int_ for 8 carboxylic acids.

**Figure 2.**
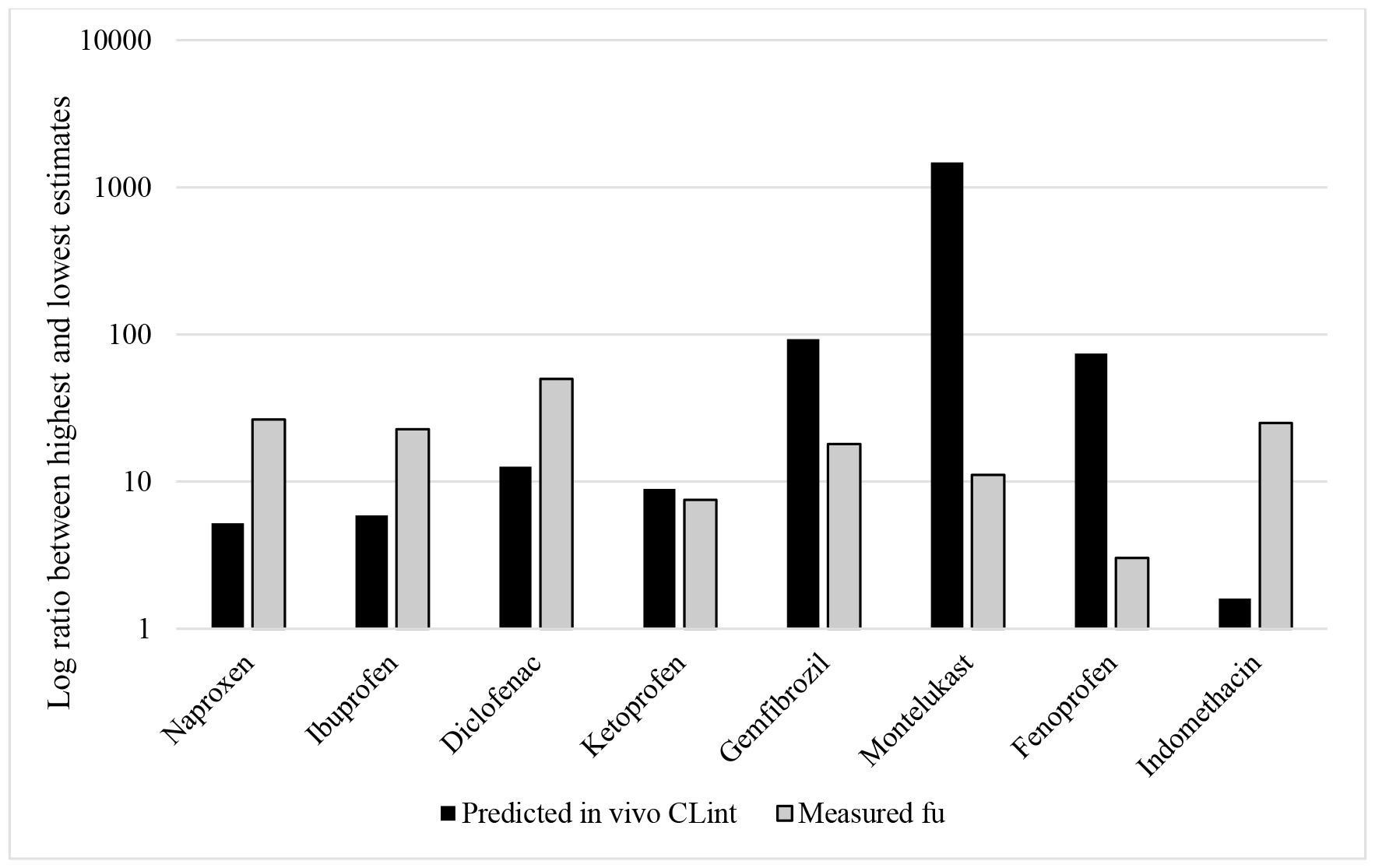
Ratios between lowest and highest reported predicted *in vivo* CL_int_ and *in vitro* f_u_ for 8 carboxylic acids.

Unfavorable human pharmacokinetics for one carboxylic acid predicted in one laboratory may be contradicted by results from another laboratory (and vice versa), such as for diclofenac (CL_H_ predicted to be low to high; 162-fold difference between lowest and highest predicted CL_H_), montelukast (CL_H_ predicted to be low to very high; 245-fold difference between lowest and highest predicted CL_H_), gemfibrozil (CL_H_ predicted to be very low to moderate; 1,083-fold difference between lowest and highest predicted CL_H_), fenoprofen (CL_H_ predicted to be very low to low; 205-fold difference between lowest and highest predicted CL_H_) and naproxen (CL_H_ predicted to be very low to low; 135-fold difference between lowest and highest predicted CL_H_).

The selection of data has a major influence on established correlations between predicted and observed CL_H_. A previous study showed that the correlation coefficient (R^2^) for predicted and observed log CL_H_ may vary between near 0 and ca 0.8-0.9 depending on the choice of *in vitro* CL_int_ and f_u_ data (Fagerholm et al. 2022).

High R^2^-values are often reported. Diclofenac, fenoprofen, gemfibrozil, ibuprofen, ketoprofen, montelukast and naproxen were included in a prediction study with human hepatocytes by Sohlenius-Sternbeck et al. 2010 at AstraZeneca. A R^2^ of 0.72 for predicted *vs* observed log *in vivo* CL_int_ was reached. A R^2^ of similar size (0.78) was reached in a hepatocyte study by Riley et al. 2005 (also at AstraZeneca) where diclofenac, fenoprofen, gemfibrozil, ibuprofen, indomethacin, ketoprofen and montelukast were included. In another study where diclofenac and montelukast were included, Riley et al. reached a R^2^ of 0.83 (McGinnity et al 2007). Fenoprofen and ketoprofen were also included in a human hepatocyte study by Yamagata et al. 2017 (Merck and ex-AstraZeneca). A R^2^ of 0.865 was reached in that study. Different f_u_-values were selected in these studies and used f_u_-values for each of the carboxylic acids were different (up to 10-fold) than the calculated averages for these. In a study by Keefer et al. 2023 (Pfizer), R^2^ was 0.63 and the *in vivo* CL_int_ and CL_H_ of montelukast were overpredicted by ca 400- and 30-fold (predicted very high CL_H_ and observed very low CL_H_), respectively.

Considerably lower R^2^-estimates (ca 0.2-0.4) were found by, for example, Stringer et al. 2010 and Fagerholm et al. 2022. Following retrospective selection of favorable f_u_-values the R^2^ doubled (Fagerholm et al. 2022). This indicates a significant difference between prospective predictions and fitted predictions.

Thus, quite large interlaboratory variability and low reproducibility was found for these selected carboxylic acids. This is believed to have a significant impact on human pharmacokinetic, exposure and dose predictions. The findings show general limitations with human hepatocytes and plasma protein measurements when predicting the pharmacokinetics of carboxylic acids.

Preferably, diclofenac, gemfibrozil and montelukast (the carboxylic acids with the largest differences between highest and lowest predicted CL_int_ • f_u_) are included in validation studies with human hepatocytes and microsomes and their mean f_u_ are used (alternatively preselected f_u_-values).

When using mean predicted values for CL_int_ and f_u_, a R^2^ of 0.42 (and positive intercept and slope<1) was found for predicted *vs* observed log CL_H_ for the 8 carboxylic acids. Retroactive selection of favorable and unfavorable values resulted in R^2^ values from 0 to 1. This clearly demonstrates the impact of data selection and subjectivism.

Unbound *in vitro* CL_int_ is estimated using unbound fraction in hepatocyte incubations. All of the selected carboxylic acids except for montelukast (which is highly bound) have been shown to have high (60 to 99 %) unbound fraction in hepatocyte incubations. Thus, interlaboratory differences of this parameter is not believed to have a great impact on the overall variability of predicted CL_int_ and CL_H_.

### Hepatocytes *vs* log D for prediction of *in vivo* CL_int_

The data set with the apparently highest number of carboxylic acids (Keefer et al. 2023) contains 39 carboxylic acids with *in vitro* hepatocyte CL_int_ and log D (both measured at the same laboratory), *in vivo* CL_int_ and *in vitro* f_u_. This data set has 748 measured *in vitro* CL_int_-values, out of which 160 (21 %) are below the limit of quantification. 18 carboxylic acids were excluded as their *in vitro* CL_int_ was below the limit of quantification.

The R^2^ for log hepatocyte predicted *in vivo* CL_int_ *vs* log *in vivo* CL_int_ was 0.34 (n=39; Figure 3). The corresponding R^2^ for log D *vs* log *in vivo* CL_int_ was 0.40 (same set of 39 carboxylic acids; Figure 4). The R^2^ between log D *vs* log *in vivo* CL_int_ for 64 carboxylic acids was 0.47.

**Figure 3.**
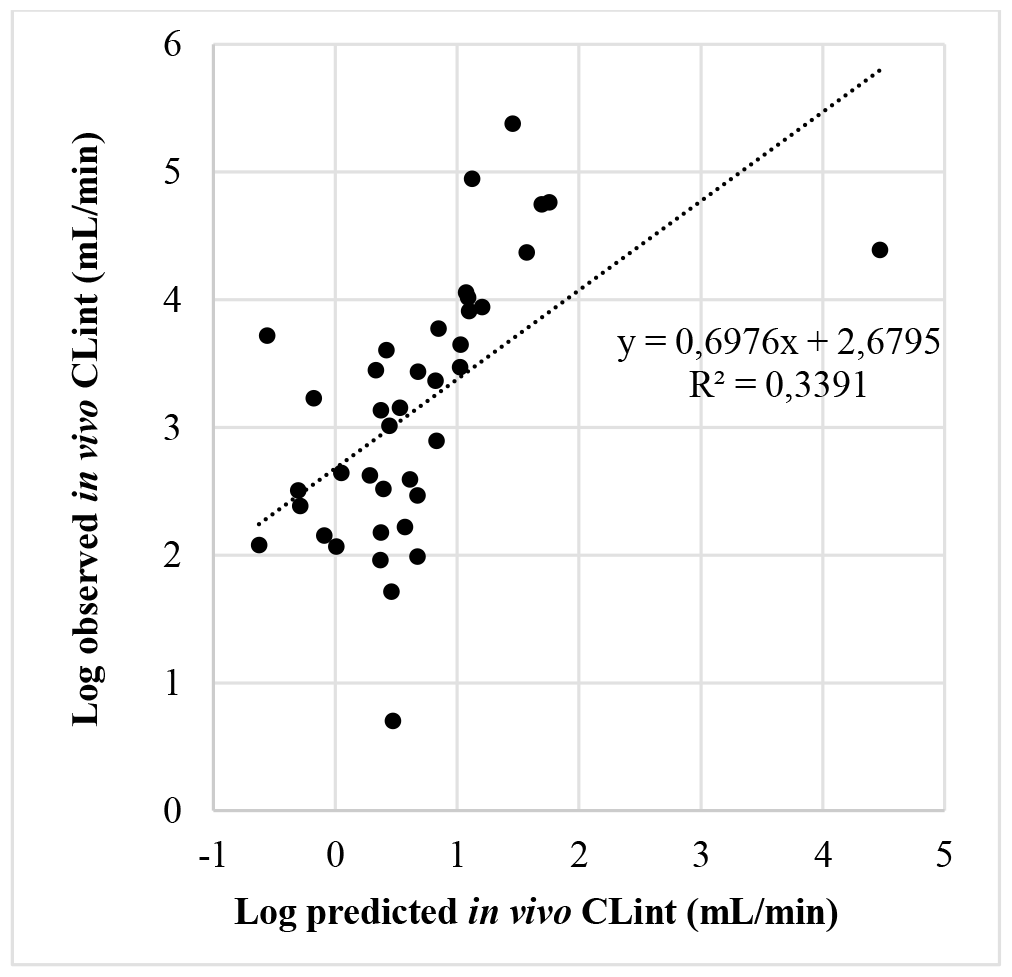
Relationship between log hepatocyte predicted *in vivo* CL_int_ and log observed *in vivo* CL_int_ for 39 carboxylic acids.

**Figure 4.**
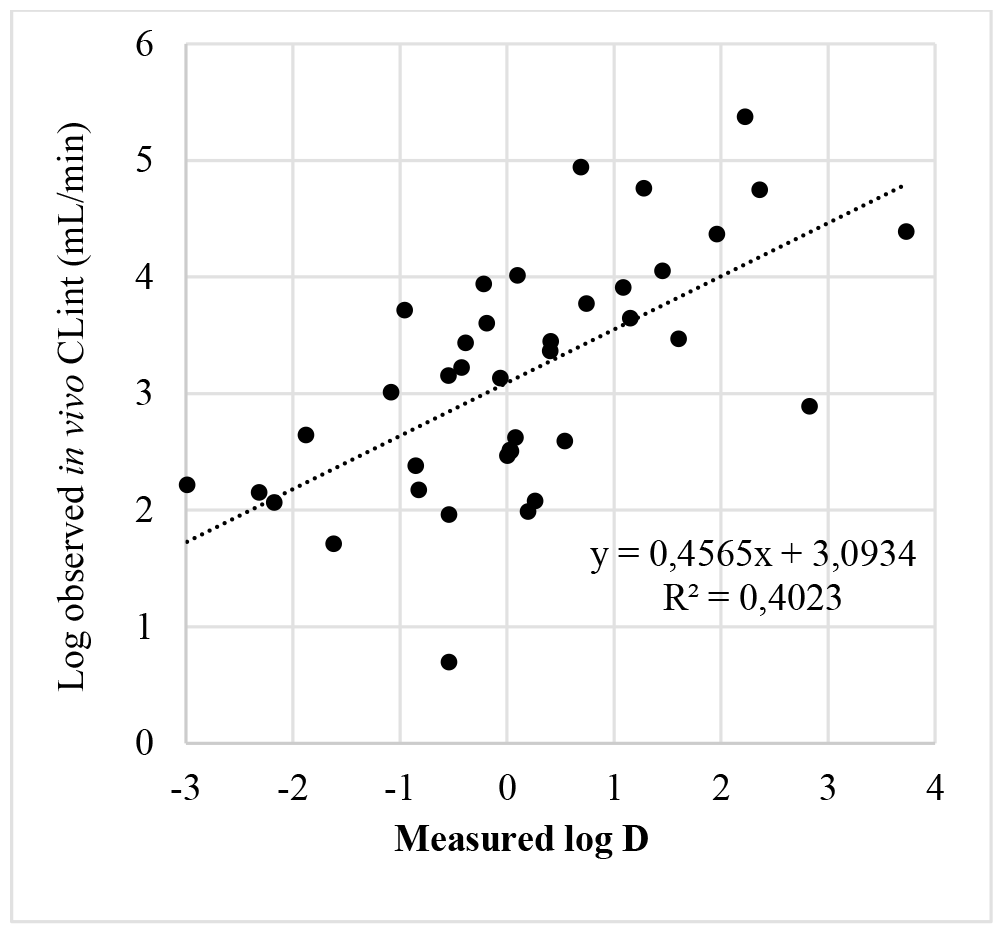
Relationship between measured log D and log observed *in vivo* CL_int_ for 39 carboxylic acids (same as in Figure 4).

With another smaller set of carboxylic acids (n=10), the R^2^ for log hepatocyte predicted *in vivo* CL_int_ *vs* log *in vivo* CL_int_ was 0.28 (Sohlenius-Sternbeck et al. 2010). Consistent with findings above, the R^2^ for log D *vs* log *in vivo* CL_int_ was higher (0.33).

Overall and for carboxylic acids, measured log D from one laboratory had a higher correlation *vs* log *in vivo* CL_int_ and wider range than human hepatocyte data from one laboratory.

It should be noted that the variability and selection of measured values could have influenced the results. The role of hepatic (vs extrahepatic) metabolism is not clear, and therefore, the degree of overestimation of hepatic CL_int_ for each carboxylic acid is unknown.

### ANDROMEDA vs hepatocytes and log D for prediction of *in vivo* CL_int_

The Q^2^ (forward-looking R^2^) between *in silico* (ANDROMEDA) predicted and measured log *in vivo* CL_int_ for 12 carboxylic acids not used in the training set of the CL_int_-model was 0.86 (Figure 5). The corresponding R^2^ (same set of carboxylic acids) for hepatocytes and log D were 0.67 and 0.66, respectively.

**Figure 5.**
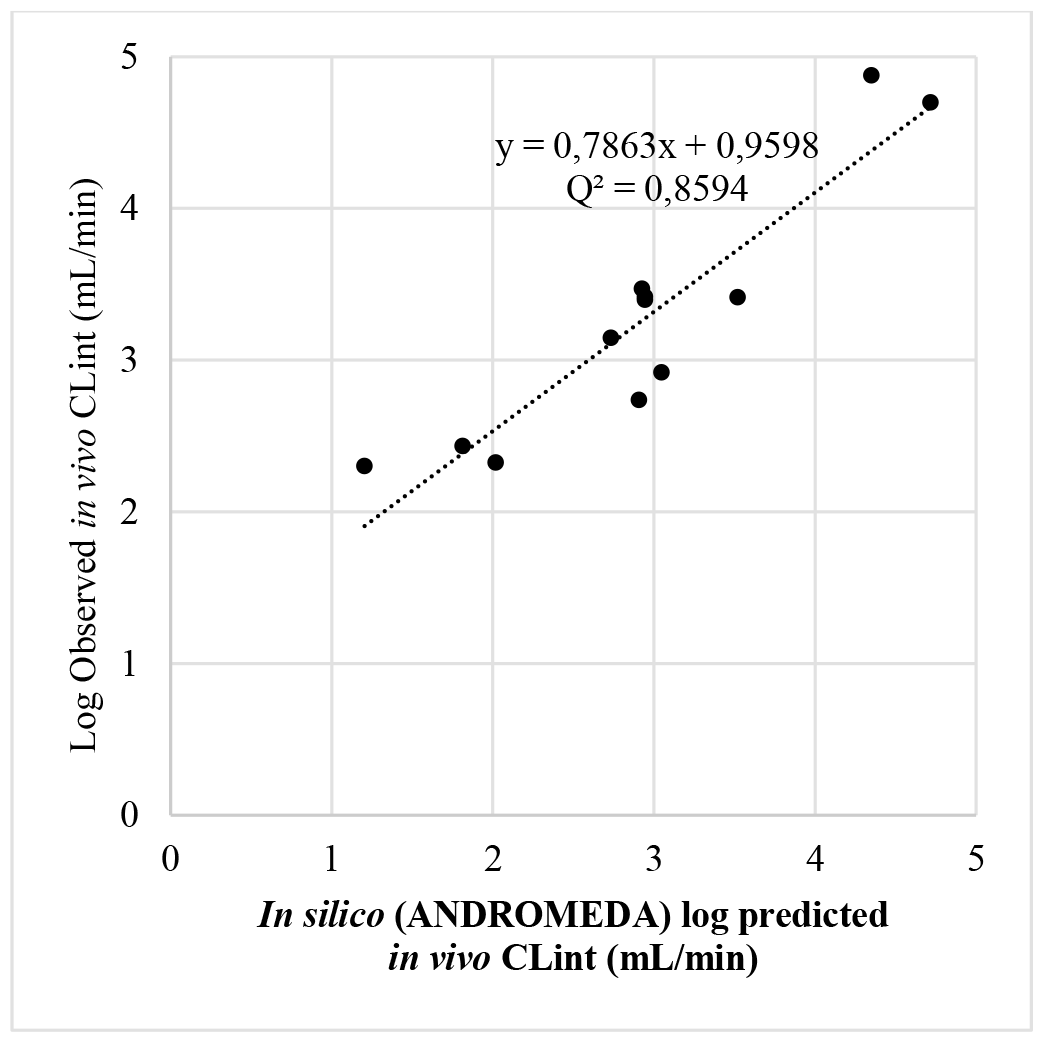
Relationship between *in silico* log predicted (ANDROMEDA by Prosilico version, 1.0) and log observed *in vivo* CL_int_ for 12 carboxylic acids.

ANDROMEDA produced a lower maximum prediction error compared to hepatocytes (12.5 *vs* 13.5-fold) and also predicted the *in vivo* CL_int_ for all carboxylic acids out of reach for the hepatocyte assay. Thus, in this comparative study it clearly outperforms human hepatocytes and log D. This is in line with a previous head-to-head study where ANDROMEDA reached a Q^2^ of 0.48 and human hepatocytes a R^2^ of 0.38 (Fagerholm et al. 2022).

Among advantages with ANDROMEDA is that it (in opposite to *in vitro* measurements) gives that same results upon repeated predictions. It also predicts the *in vivo* CL_int_ and CL_H_ of diclofenac, fenoprofen, gemfibrozil, montelukast and naproxen (which demonstrate very large interlaboratory variability) well.

The CL_int_ model in ANDROMEDA version 2 has 22 % higher relative Q^2^ and 8 % lower relative RMSE compared to version 1, which indicates that even better performance is expected for carboxylic acids with an updated version.

## Conclusion

Very large interlaboratory variability was demonstrated for plasma protein binding and hepatocyte assays. Log D, and especially ANDROMEDA, outperformed the hepatocyte assay for the prediction of CL_int_ of carboxylic acids *in vivo* in man.

